# A reference genome and its epigenetic landscape of potential *Orychophragmus violaceus*, an industrial crop species

**DOI:** 10.1101/2023.09.21.558835

**Authors:** Changfu Jia, Yukang Hou, Qiang Lai, Yuling Zhang, Rui Wang, Jianquan Liu, Jing Wang

## Abstract

*Orychophragmus violaceus*, also called ‘er-yue-lan’ in China, is an annual plant of the family of Brassicaceae. The seed oil of *O. violaceus* contained two specific di-hydroxy fatty acids which were produced by functional divergence between two *FAD2* WGD copies determine its industrial property. Here, we assembled a high-quality chromosome-level genome of *O. violaceus* via based on PacBio CLR technology and its cis-regulatory landscape of five mature tissues, including root, leaf, flower, seed and stem, based on ATAC-seq technology. 1.2 Gb draft genomic sequences were anchored to 12 pseudo-chromosomes and 49904 protein-coding genes were annotated on these chromosomes. To fully understand the epigenetic landscape of *O. violaceus*, we further performed WGBS-seq for leaf, flower and silique tissues. In total, our multi-omics data provide opportunity to find out the differences between two WGD copies and also a valuable resource for downstream breeding effort of this potential industrial species.

## Introductions

The seed oil of *<u>Orychophragmus</u> violaceus* enriched in lineage-specific di-hydroxy fatty acids (nebraskanic fatty acid, 7,18-OH-24:1Δ15; wuhanic fatty acid, 7,18-OH-24:2Δ15,21)[1]. These fatty acids give the seed oil of *O. violaceus* higher temperature-stability and its seed oil was improved have great industrial properties[2]. Previous studies also confirmed a lineage-specific whole genome duplication (WGD) and specific transposable element explosion occurred in *O. violaceus*, highlighting the interesting evolutionary process[3-5]. ATAC-seq were served as robust approach to detect whole-genomic wide cis-regulatory landscape.[6, 7] The active transcriptional regions always have lower CHG and CHH methylation level[8]. In order to unveil the regulatory landscape of *O. violaceus*, we generated a high-quality chromosome-level genome and combined ATAC-seq and WGBS-seq with multi-tissues.

## Results

### Genome assembly

We firstly performed a survey for *O*.*violaceus* genome to obtain the background information. Survey estimated around 1.3Gb genome size of *O*.*violaceus* with almost 2.0% heterozygous rates, indicating its highly repetitive and large genome feature. Around 142x sequence depth with 179Gb PacBio CLR sequences together with 204 Gb Hi-C data and 67Gb resequencing data were obtained for assembling genome of *O. violaceus*. To accurately assemble the genome of *O. violaceus*, we got 2.3Gb genome with applying polyploid parameter of CANU [9]which preferred separate the heterozygous regions into two independent sequences, which was advised for reducing the error rates in generating genome assembly of highly repetitive large genome. After deduplication, 1.3 Gb size were gotten and around 1.2 Gb sequences were anchored to 12 pseudo-chromosomes. The completeness and accuracy of the assembled genome were validated using benchmarking universal single-copy orthologues (BUSCO) showed that 98.6% complete plant orthologues were recalled. We further calculated the contigN50 of our genome with 2.75 Mb. Collectively, we generated a high-quality reference genome of *O. violaceus* via PacBio CLR technology (Table 1).

**Table 1.** Statistics of assembled contigs for *O. violaceus*.

| **PacBio Sequel sequencing** |  |
| --- | --- |
| Raw data(Gb) | 179 |
| Sequencing depeth(×) | 142 |
| Average reads length (bp) | 21275 |
| Reads N50 (bp) | 30258 |
| **Hi-C sequencing** |  |
| Clean data(Gb) | 204 |
| Sequencing depeth(×) | 157 |

| **Monoploid genome assembly and annotation** |  |
| --- | --- |
| Estimated genome size (Gb) per 1 C | 1.3 |
| Contig Assembly size(Gb) | 1.28 |
| Scaffold Assembly size(Gb) | 1.19 |
| Contig N50(Mb) | 2.75 |
| BUSCO completeness of assembly (%) | 98.6 |

### Genome annotation

The majority of the genome of *O. violaceus* were occupied by TE elements with 75.47% of genomes in total. Specifically, these TE were classified into family level and LTR/Copia, LTR/Gypsy and LTR/unknown were the most popular TE family in *O. violaceus* genome (Table 2). A total of 49904 protein coding genes were annotated in our genome with 3894 bp average gene length, 5.19 average exon numbers per gene and 1046 average CDS length (Table 3). Gene annotation of BUSCO indicates the goodness of our genome assembly (Table 4). The genomic features of our genome showed that *O. violaceus* have undergone a lineage specific TE expansion event, which is consistence with other genome assemblies of *O. violaceus*.

**Table 2.** TE content of *O. violaceus* genome.

| TE type | Total size (bp) | Percentage of genome (%) |
| --- | --- | --- |
| **Class I: Retrotransposon** | 682,147,775 | 53.06% |
| **LTR Retretransposon** | 680,867,308 | 52.96% |
| Copia | 291,457,532 | 22.67% |
| Gypsy | 238,617,482 | 18.56% |
| unknown | 150,792,294 | 11.73% |
| **LINE** | 1,280,467 | 0.10% |
| **Class II: DNA transposon** | 248,772,353 | 19.35% |
| Helitron | 141,807,979 | 11.03% |
| CACTA | 54,313,799 | 4.23% |
| Mutator | 31,914,567 | 2.48% |
| PIF_Harbinger | 9,380,070 | 0.73% |
| Tcl_Mariner | 1,725,727 | 0.13% |
| hAT | 9,630,211 | 0.75% |
| **Unclassified** | 39,116,002 | 3.04% |
| **Total** | 970,043,238 | 75.47% |

**Table 3.** Gene features of *O. violaceus* genome.

| Feature | number/length |
| --- | --- |
| Gene number in Chr | 49,904 |
| Average gene length | 3,894 |
| Average exon number per gene | 5.19 |
| Average CDS length | 1,046 |

**Table 4.** BUSCO of gene annotation of *O. violaceus* genome.

| BUSCO | Gene-Number | Gene-Percentage |
| --- | --- | --- |
| Complete BUSCOs (C) | 1539 | 95.40% |
| Complete and single-copy BUSCOs (S) | 1005 | 62.30% |
| Complete and duplicated BUSCOs (D) | 534 | 33.10% |
| Fragmented BUSCOs (F) | 12 | 0.70% |
| Missing BUSCOs (M) | 63 | 3.90% |
| Total BUSCO groups searched | 1614 | 100.00% |

### Epigenetic features of *O. violaceus* genome

We first determined the genomic features of *O. violaceus* genome (Fig. 1A). Defining the genomic region with highest Tn5-occupy frequency as accessible chromatin regions (ACRs), we identified 31,389, 25,467, 27,251, 30,773, 29,690 ACRs in flower, leaf, root, seed and stem, respectively. The ACRs were normalized around the transcriptional start site (TSS) and transcriptional end site (TES). TSS and TES showed highest ACR frequencies and its flanking sequences were gradually decreased from its center (Fig. 1B). We identified CpG, CHG and CHH across the *O. violaceus* genome. Considering the high TE explosion of *O. violaceus* genome which might interrupt the gene features, we divided annotated genes into gene-with-TE and gene-without-TE. The methylation levels of overall genes among CpG, CHG and CHH are obviously higher than the genes without any TEs. The methylation level often links to the gene expression level. These results indicates that the TE insertion event could largely change the methylation level of protein-coding genes (Fig. 1C).

**Fig. 1.**
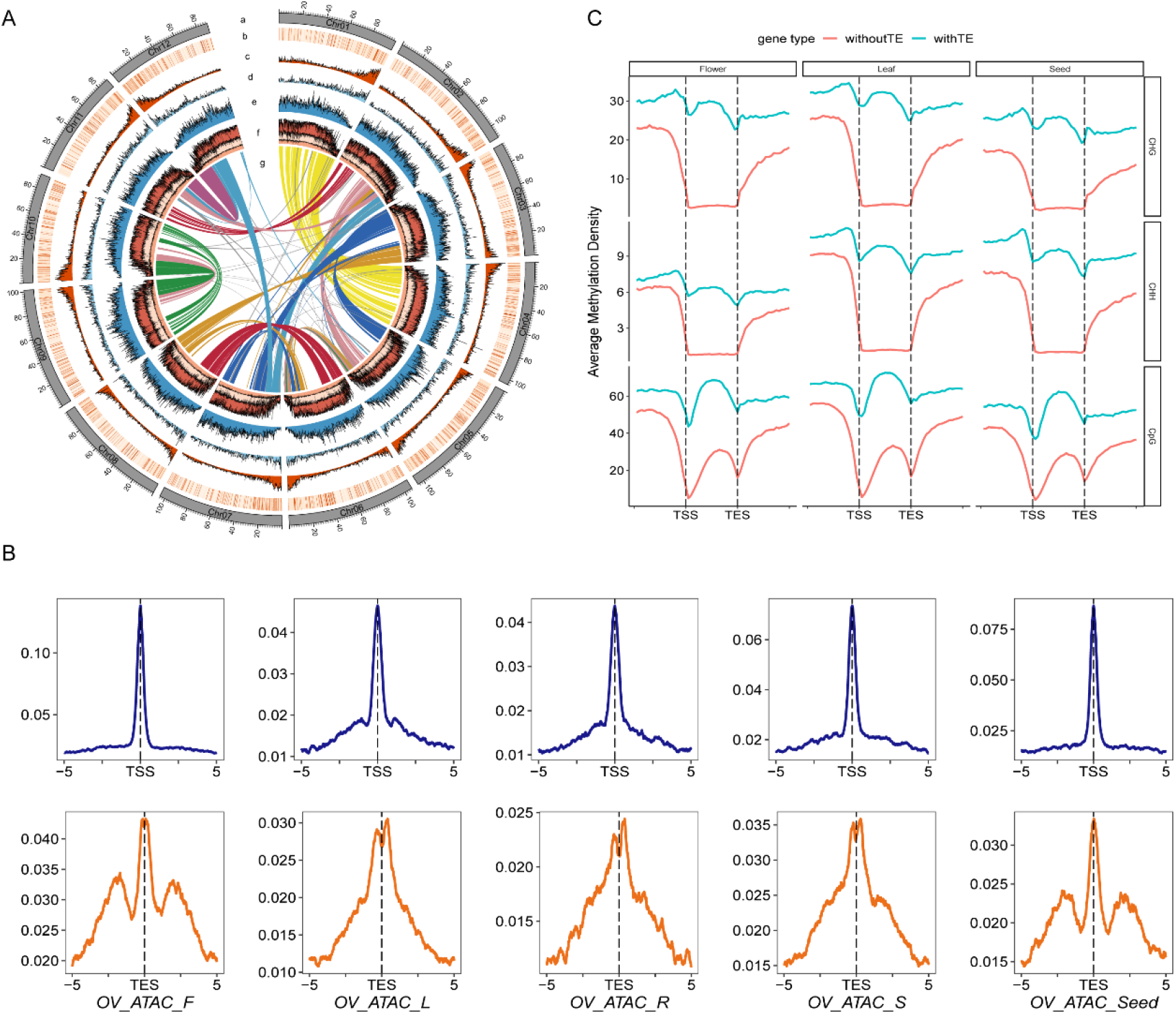
A) Genomic features of the O. violaceus genome. (a) The 12 chromosomes in the genome of O. violaceus; (b) Gene expression density in leaf; (c) Gene density; (d) ACR density distribution in leaf; (e) LTR/Copia density distribution; (f) From high to low is the methylation density of CG, CHG, CHH (g) Syntenic blocks in O. violaceus genome. B: The enrichment of ACR form 5kb downstream of transcriptional start sits (TSS) and transcriptional end sites (TES) across five tissues. C) The differences between genes with/without TE of CpG, CHG and CHH methylation levels in flower, leaf and seed tissues.

## Conclusion

In this study, we de-novo assembled a high-quality reference genome of *O. violaceus*. Combining multi-omics data in several tissues, we dissected the epigenetic landscape and highlighted methylation changes caused by the abundant TE insertion events specific to *O. violaceus*. Our data sets provide a valuable resource and further study is needed to integrate these data with additional RNA-seq data to investigate the functional divergence and the influence of epigenetic changes of WGD gene pairs of *O. violaceus*.

## Materials and Methods

### Plant materials and genome sequencing

The Genome Sequencing of *O. violaceus* and the multi-tissue transcriptome materials for gene annotation were collected in Chengdu City, Sichuan Province (30°37’45.4”N, 104°05’32.5”E). The extracted DNA was used for PacBio CLR sequencing and second-generation sequencing. For the third-generation sequencing, DNA was firstly interrupted with g-TUBE and then hairpin structure adapters are added to both ends of the DNA. BluePippin is used to screen the target fragments. Finally, the high-quality purified DNA is used. SMRTbell Templat Prep Kit 1.0 constructs a PCR-free SMRTbell library, which is finally sequenced on the PacBio Sequel II platform. A total of 179Gb data with a depth of about 142x was measured, and the N50 was 30,258bp. The results of third-generation sequencing were mainly used for the assembly of the *O. violaceus* reference genome. For the short-read sequencing, Illumina’s Genomic DNA Sample Preparation kit is used to construct a paired-end Illumina DNA sequencing library with an insert library length of 400bp, and the Illumina HiSeq 2500 platform is used for sequencing. A total of 67Gb data with a depth of approximately 54x is obtained. This data is used for Genome survey evaluation and genome assembly error correction. High-throughput chromosome conformation capture (Hi-C) sequencing has been widely used in the mounting step of genome assembly. We selected 3g fresh leaves for liquid nitrogen grinding, cell cross-linking, and DpnII endonuclease digestion, end-repaired, labeled with biotin, and linked to obtain blunt-end fragments. After purification, the DNA was randomly sheared to 300-500bp fragments. After quality control, the library was sequenced using the Illumina Novaseq 6000 platform to sequence the 150bp paired-end reads and 203Gb clean reads were obtained.

### Genome assembly and quality assessment

We firstly used Illumina short reads to estimate the genome size of *O. violaceus* using GenomeScope2 with default parameters[10]. Based on the estimated heterozygous rates and around 1.3Gb genome size, we conducted CANUv2.2 [9]to assembled the Pacbio CLR reads with specific parameters’ batOptions=-dg 3 -db 3 -dr 1 -ca 500 -cp 50’ which was advised for generating high heterozygous and repetitive genomes. 2.13Gb genome sequences were obtained by CANU. NEXTDENOVO was used to reduce error rates of the above assembly[11]. Purge_haplotigs v1.1.1[12] was performed to deduplicates. We finally anchored the purged genome assembly with around 1.3Gb to 12 chromosomes via 3dDNA software and manually adjusted by juicer[13]. Genome sequences were submitted to BUSCO v4.0.5 [14]with embryophyta_odb10 download at 16-Oct-2020 as reference database to test the quality of our genome assembly.

### Gene prediction and functional annotation

We firstly predicted the transposable elements of *O. violaceus* genome using EDTA v1.9.3 pipeline[15]. For gene prediction, we combined three strategies including homology-based prediction, transcriptome-based prediction, and *ab initio* prediction. For homologous-based prediction, protein coding genes of six selected species, including *Arabidopsis thaliana, Populus trichocarpa, Vitis vinifera, Brassica napus, Brassica rapa and A*.*lyrata* were aligned to our reference genome using TBLASTN[16]with 1e-5 as cutoff. The complete gene structure were further confirmed via Genewise v2.4.1[17]. For transcriptome-based prediction, we firstly trimmed adapters and filtered low-quality reads via fastp with default parameters[18]. The trimmed RNA-seq reads were aligned to our genome using hisat2 with default parameters[19]. We next conducted genome-free and genome-guided method to get assembled transcripts using Trinity v2.8.4[20]. The obtained transcripts were aligned to the genome assembly by both BLAT[21] and gmap[22] under the PASA pipeline[23] to get mRNA-based gene prediction evidence. For *ab initio* prediction, we incorporate the homologous and transcriptomic based gene models to train gene models using Augustus[24]. All the gene models predicted by these three strategies were sent to EvidenceModeler v1.1.1[23] with different weight to obtain integrative gene models. Finally, based on the mRNA-seq data, the integrative gene models were updated three rounds for adding alternative splicing event and accurately boundary of coding regions using PASA pipeline[23].

For the functional annotation, we firstly aligned our predicted gene models to well-known public databases using blastp with 1e-5 as cutoff[16], including the protein families database (Pfam)[25], the NCBI non-redundant protein database (NR)[26] and the Swiss-Prot protein database[27]. GO and KEGG terms were annotated by InterProScan[28].

